# Multiplexed temporally focused light shaping for high-resolution multi-cell targeting

**DOI:** 10.1101/216135

**Authors:** Nicolò Accanto, Dimitrii Tanese, Emiliano Ronzitti, Clément Molinier, Zachary L. Newman, Claire Wyart, Ehud Isacoff, Eirini Papagiakoumou, Valentina Emiliani

## Abstract

Patterning light at the single-cell level over multiple neurons in the brain is crucial for optogenetic photostimulation that can recapitulate natural activity patterns and, thereby, determine the role of specific components of brain activity in behavior. To this end we have developed a method for projecting three-dimensional, 2-photon excitation patterns that are confined to many individual neurons. The new versatile optical scheme generates multiple extended excitation spots in a large volume with micrometric lateral and axial resolution. Two-dimensional temporally focused shapes are multiplexed several times over selected positions, thanks to the precise spatial phase modulation of the pulsed beam. This permits, under multiple configurations, the generation of tens of axially confined spots in an extended volume, spanning a range in depth of up to 500 μm. We demonstrate the potential of the approach by performing multi-cell volumetric excitation of photoactivatable GCaMP in the central nervous system of Drosophila larvae, a challenging structure with densely arrayed and small diameter neurons, and by photoconverting the fluorescent protein Kaede in zebrafish larvae. Our technique paves the way for the optogenetic manipulation of a large number of neurons in intact circuits.

## Introduction

Nowadays, the availability of a growing number of optogenetic actuators^1,2^, calcium^3^ or voltage reporters^4^ and photoactivable proteins^5–9^, enables the use of light to control and monitor neuronal activity, visualize cell morphology and track neuronal connectivity in intact brain preparations^10–15^. At the same time, the investigation of complex brain circuits has spurred the development of new optical approaches for fast imaging and manipulation of neurons in large volumes.

In the simple cases of sparse genetic expression, low scattering tissues or superficial targets, fast volumetric imaging can be achieved under (1P) illumination^16–18^. However, two-photon (2P) illumination is favourable to reach deeper regions in scattering tissues. 2P volumetric imaging was achieved either with scanning approaches, by quickly scanning single or multiple beams over planes or objects of interests^19–23^, or with parallel approaches, based on the simultaneous illumination of multiple targets in a volume and on extended depth of field detection^24,25^.

Similarly, fast volumetric neuronal activation can be achieved using relatively simple illumination methods based on wide field 1P illumination^5,26,27^. However, reaching in-depth single-cell resolution requires replacing 1P with 2P excitation (2PE), and calls for more sophisticated illumination approaches. Efficient 2PE of multiple channels or molecules expressed on a cell membrane can be achieved by raster or spiral scanning a diffraction-limited spot^28,29^ over the cell soma. Alternatively, the entire cell surface can be covered at once using scan-less parallel light shaping approaches such as low-numerical aperture (NA) Gaussian beams, Computer-Generated Holography (CGH)^30–33^ or the Generalized Phase Contrast (GPC) method^34,35^. In the case of parallel illumination, the loss of axial resolution consequent to the increase of the lateral spot dimension can be compensated for by the use of temporal focusing (TF)^36,37^. TF uses a diffraction grating placed at a plane conjugated with the objective focal plane. By dispersing the spectral frequencies of ultra-short excitation pulses the TF-grating temporally smears the pulse away from the focal plane, which remains the only region irradiated at peak powers efficient for 2PE, thus reaching micrometer-axial resolution independently of the lateral spot size. Remarkably, TF is also critical for preserving the excitation patterns in scattering tissues^33,35,38^.

Parallel approaches present the great advantage of minimizing the illumination time with respect to their scanning counterparts. This can be seen as follows: the temporal resolution for scanning illumination, TR,scan, roughly equals the illumination time per spot (tdwell) multiplied by the number of scanned positions and by the number of targets, while that for parallel approaches, TR,paral, is only given by tdwell. As a consequence, for volumetric multi-cell targeting, TR,scan can strongly exceed the value of TR,paral and 3D parallel illumination remains the only option to achieve millisecond temporal resolution^39^.

Although not yet fully parallel, a first implementation of 3D multi-target illumination used CGH to generate multiple diffraction-limited spots^40–42^ at the positions of the targeted cells and scanned all of them simultaneously over the cell membranes using a galvanometric system^12,43^. Combined with optogenetics, scanning-CGH proved the capability to target multiple cells. Yet, the need of scanning over the cell body limited the achievable temporal resolution (>20 ms for single AP generation) and precision (temporal jitter >6ms)^12,44^. Moreover using focused light at saturation power to compensate for the small spot surface generated important out-of-focus excitation^28,39,45^.

Full parallel illumination of multiple targets with minimal temporal resolution was achieved by generating extended holographic shapes in 3D^13,46,19^. CGH combined with optogenetics, enabled *in vivo* and in *vitro* action potential (AP) generation with <3 ms temporal resolution and <1 ms temporal jitter^47–49^. However, a major issue related to 3D-CGH in its conventional configuration is its incompatibility with the use of TF, which requires the focalization of 2D holographic patterns at the TF grating. Without TF, the axial resolution of extended holographic spots deteriorates, especially in the case of spatially close targets, thereby limiting 3D-CGH to the illumination of sparsely distributed targets.

We recently overcame this issue by using two spatial light modulators (SLMs), the second one placed after the TF-grating^13^ and by tiling the SLMs to form multiple co-aligned holograms, which independently controlled the lateral shape and position (SLM1) and the axial position (SLM2) of each pattern. This strategy enabled the remote axial displacement of a temporally focused shape as well as focalizing different temporally focused patterns at axially distinct planes. Yet, the necessity of tiling the SLMs physically limited the number of planes that could be addressed. Moreover, this approach was hardly compatible with other beam-shaping techniques.

Here we demonstrate a novel configuration for multiplexed temporally focused (MTF) light shaping, which removes all these limitations. The main concept behind it is to decouple the light shaping into two completely independent steps: a first beam-shaping unit generates and focuses a desired 2D shape on the TF grating; in a second step a SLM, placed after the grating, axially and laterally multiplexes the 2D shape creating a 3D distribution of an arbitrary number of replicas. A similar multiplexing scheme was recently used to replicate GPC shapes in 1P regime^50^, or temporally focused low-NA Gaussian beams^51^. In the former case, the lack of 2PE and TF limited the achievable axial resolution, whereas in the latter the method was restricted to the generation of a static and single-size Gaussian spot.

In this work instead, we set up a fully flexible scheme, easily compatible with different light shaping approaches, such as CGH and GPC, for the 3D multiplexing of extended shapes with high axial resolution. We validate the spatial precision of MTF light shaping by performing in vivo multi-cell volumetric excitation of superfolder photoactivatable GCaMP (sPA-GCaMP)^6^ in fruit flies, and Kaede photoconversion^52^ in zebrafish larvae. In both scenarios, we show the specific targeting of individual neurons selected out of dense arrays and distributed over multiple planes.

## Optical system

The optical system for multiplexed temporally focused light shaping, schematically represented in **Fig. 1a**, consisted of three main parts: (i) the beam-shaping unit, which generated a 2D illumination pattern, described at the objective focal plane by the function *f(X,Y)*; (ii) the temporal focusing unit comprising the grating and the appropriate lenses, which temporally focused the 2D pattern at the objective focal plane; (iii) the multiplexing unit, constituted of a liquid-crystal SLM (SLM2), which created, using CGH, the desired 3D distribution of diffraction-limited spots at the sample positions *Xi*, *Yi* and *Zi*, thereby replicating the original 2D pattern *f(X,Y)* at such positions (**Fig. 1b** and Methods).

**Figure 1:**
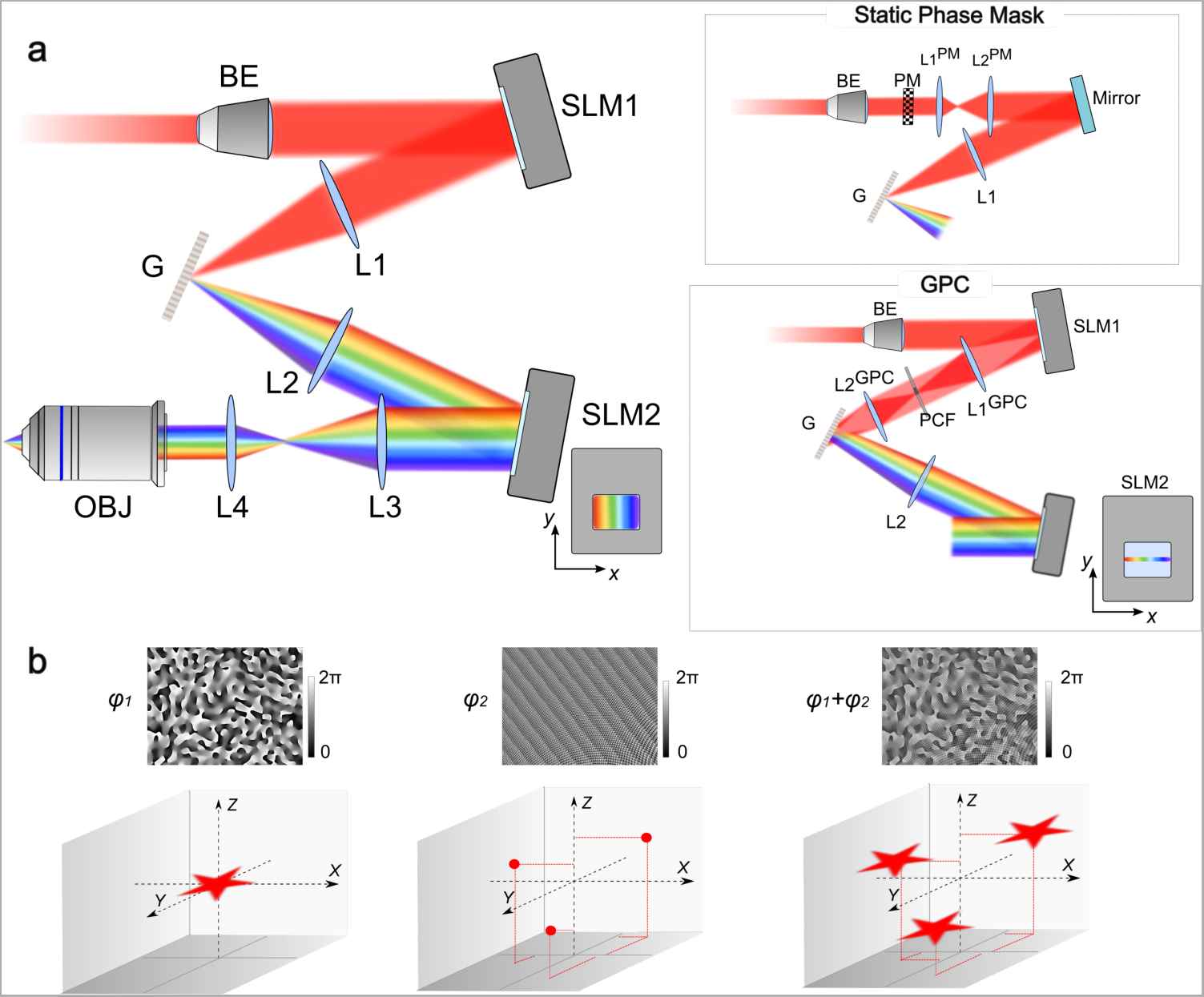
Experimental setup and principle of 3D multiplexed temporally focused excitation. (**a**) The experimental setup comprised the beam-shaping unit, the temporal focusing unit and the multiplexing unit. In a first realization, the beam-shaping unit consisted of a dynamic CGH system, composed of a beam expander (BE) to match the active area of a LC-SLM (SLM1), which performed the appropriate phase modulation. The 2D illumination pattern was then focused on the grating (G) for TF through lens L1. The first diffraction order was collimated by lens L2 and directed into the second LC-SLM (SLM2), which created the desired 3D distribution of diffraction-limited spots at the sample positions thereby replicating the 2D pattern generated by SLM1. Lenses L3 and L4 conjugated the SLM2-plane at the objective (OBJ) back focal plane and scaled the beam at the objective back aperture. Close to SLM2 it is shown the schematic of the *x*-*y* light distribution on SLM2. In a second realization, the 2D illumination pattern was a circular 20-μm diameter holographic spot created via phase modulation by a static phase mask (Inset, top). The telescope constituted by lenses L1^PM^ and L2^PM^ magnified the size of the static phase mask, to match the size of SLM2. In a third realization, the beam shaper was a GPC-interferometer (Inset, bottom). Binary phase modulation for a circular spot of 12-μm diameter was addressed to SLM1, and lens L1^GPC^ focused the beam on the phase contrast filter (PCF), which introduced a λ/2 phase retardance to the low-spatial-frequency components over the high-spatial frequencies. Finally, the lens L2^GPC^ recombined high and low-spatial frequencies to form the interference pattern at the output plane of the GPC-shaper, which coincided with the grating plane. Close to SLM2 it is shown the schematic of the *x*-*y* light distribution on SLM2: in this configuration, the dispersed beam after the diffraction grating was focused on SLM2, resulting in the illumination of only a small portion of SLM2 in the *y*-direction while the dispersion of spectral frequencies enabled to cover the LC area along the *x*-direction. (**b**) Schematic principle of 3D spatial beam multiplexing for the case of CGH. A user-defined 2D illumination pattern is generated (left panel) by addressing SLM1 with the corresponding holographic phase *φ1*. At the same time, SLM2 is addressed with a phase hologram, *φ2*, encoding diffraction-limited spots at different positions, *i*, in the 3D space (middle panel). The combination of the two holograms, φ_1_ + φ_2_, at the objective back aperture enables to generate several replicas of the original pattern at the positions *i* (right panel).

In this work, we demonstrate the 3D generation of multiple temporally focused spots using three different configurations for the beam-shaping unit: 1) CGH based on the use of a liquid-crystal SLM (**Fig. 1a**), which constituted the reference approach that we subsequently used in two different biological applications; 2) CGH using a static holographic phase mask (**Fig. 1a**, inset top); 3) GPC interferometry (**Fig. 1a**, inset bottom).

### Multiplexed Temporally Focused Computer-Generated Holography (MTF-CGH)

We first proved the strength of our system by creating 50 multiplexed temporally focused computer-generated holographic (MTF-CGH) replicas of a circular 15-μm-diameter shape using a conventional scheme for CGH based on a liquid-crystal SLM (SLM1; **Fig. 1a**). The 50 spots were visualized by measuring the 2PE fluorescence from a thin (sub-µm) rhodamine layer in a two-photon microscope equipped with a 40x, 0.8 NA objective (**Fig. 2a**). **Fig. 2b** shows three replicas of the excitation spot at three different axial planes (-250, 0, 240 μm from the focal plane). One can notice the speckled intensity profile, typical of holographic spots^53^, as well as the similarity of the speckle distribution for spots lying on different planes, confirming that each spot was indeed a replica of the original 2D shape.

**Figure 2:**
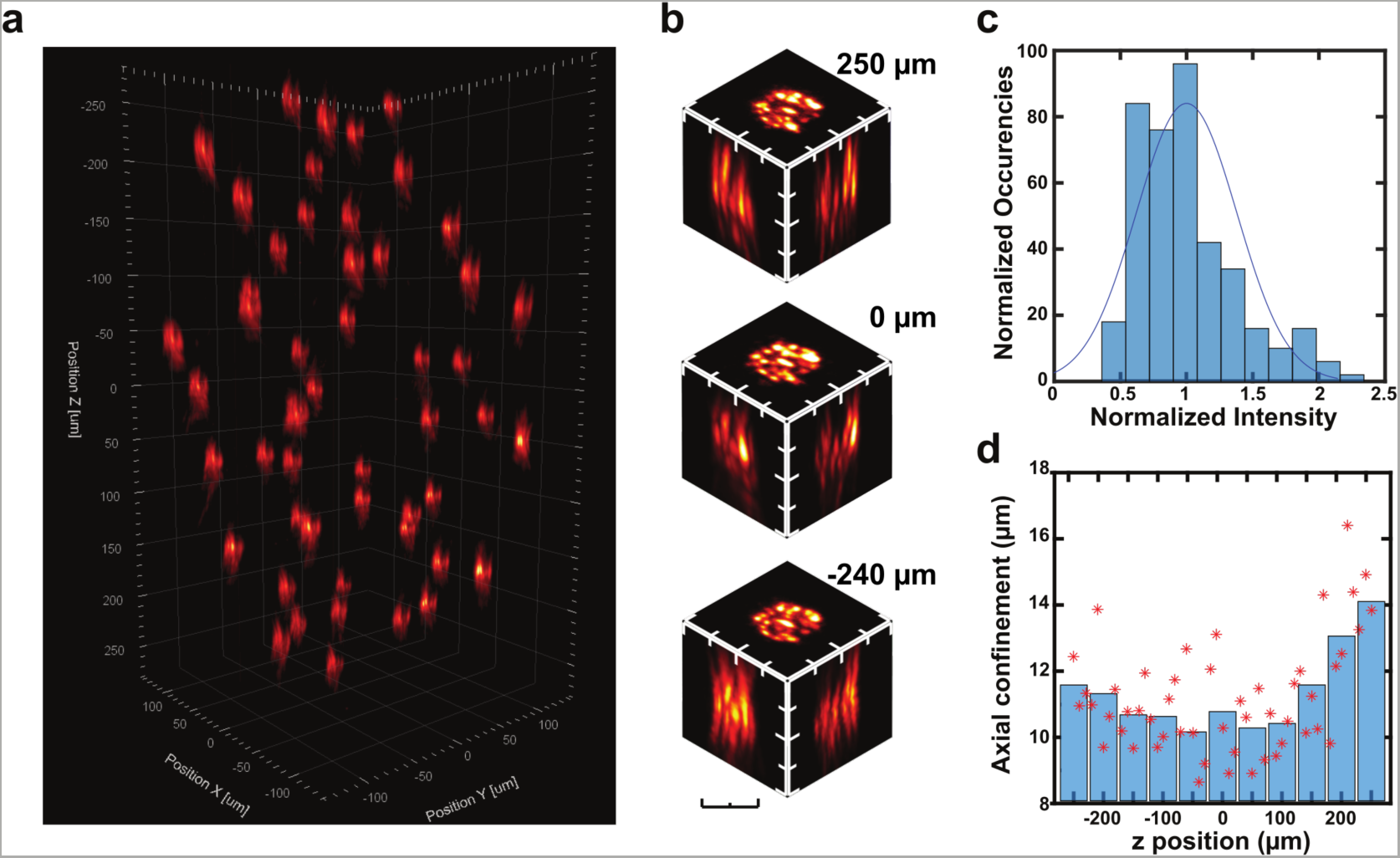
MTF-CGH with two SLMs. (**a**) 2PE fluorescence volume representation of 50 holographic circular spots of 15-μm diameter, each of them lying on a different plane, in a volume of 300×300×500 μm^3^. To record the stack from which we reconstructed the 3D volume, we used 450 mW average laser power at the back aperture of the objective (40x, 0.8 NA), by integrating 1 s per plane with the imaging camera. (**b**) *x*-*y*, *x*-*z* and *y*-*z* projections of three spots, located at *z*=-250, 0, 240 μm from the focal plane. Scale bar: 15 μm. (**c**) Histogram of the maximal 2PE fluorescence intensity for each spot, normalized to the average intensity of all spots, after diffraction efficiency correction. The results represent an average for each plane from 4 different realizations of 50-spots light configuration. (**d**) Axial confinement, calculated as the FWHM of the axial intensity profile of each spot, as a function of the *z* position. Red stars represent the individual measurements for each spot (average on 4 different realizations of 50 spots) and blue bars show the mean values in a range of 50 μm around the designated *z* position. The mean value across the whole FOE was 11.1±1.8 µm FWHM.

As previously described^13,54,55^, by calculating the phase that we sent to SLM2 with a weighted Gerchberg-Saxton algorithm we could compensate for diffraction efficiency-induced intensity variations. In this way the 2PE fluorescence from the 50 spots varied less than 40% around the mean value, corresponding to less than 25% variation in the illumination intensity (**Fig. 2c)**.

In **Fig. 2d** we plot the full-width at half maximum (FWHM) of the axial intensity distribution for the 50 spots, which was found to vary between 8 and 16 µm, with a mean value across the whole investigated field of excitation (FOE ≈300×300×500 µm^3^) of 11.1±1.8 µm and only few spots, at the very edge of the FOE in the *z* direction, reaching a FWHM >15 µm. As shown in detail in **Supplementary Fig. 1-3**, the FWHM was as small as 7 µm at the centre of the FOE and increased both in the *x*-*y* plane (~0.4% and 0.5% per µm for the *x* and *y* direction, respectively) and, as previously observed^56^, in the *z*-direction (~0.4% per µm) as we moved away from the centre of the FOE. Two main effects could cause such axial resolution broadening. First, large axial shifts required beams highly converging or diverging at the back aperture of the objective, whereas large lateral shifts corresponded to strongly tilted beams after SLM2. Therefore, in both cases, the optical elements that the beam propagated through after SLM2 introduced strong aberrations^56^ and a possible cropping of the beam with consequent loss of spectral frequencies. A second possible contribution to the axial broadening may come from spatiotemporal coupling effects related to the spatial separation of the spectral frequencies at the position of SLM2 (**Supplementary Fig. 1-3**). This latter effect could also induce the few-micrometre axial shift of the spatiotemporal focal plane when shifting the illumination spots in the direction of TF (**Supplementary Fig. 2**) not observed in the non-TF direction (**Supplementary Fig. 3**).

**Figure 3:**
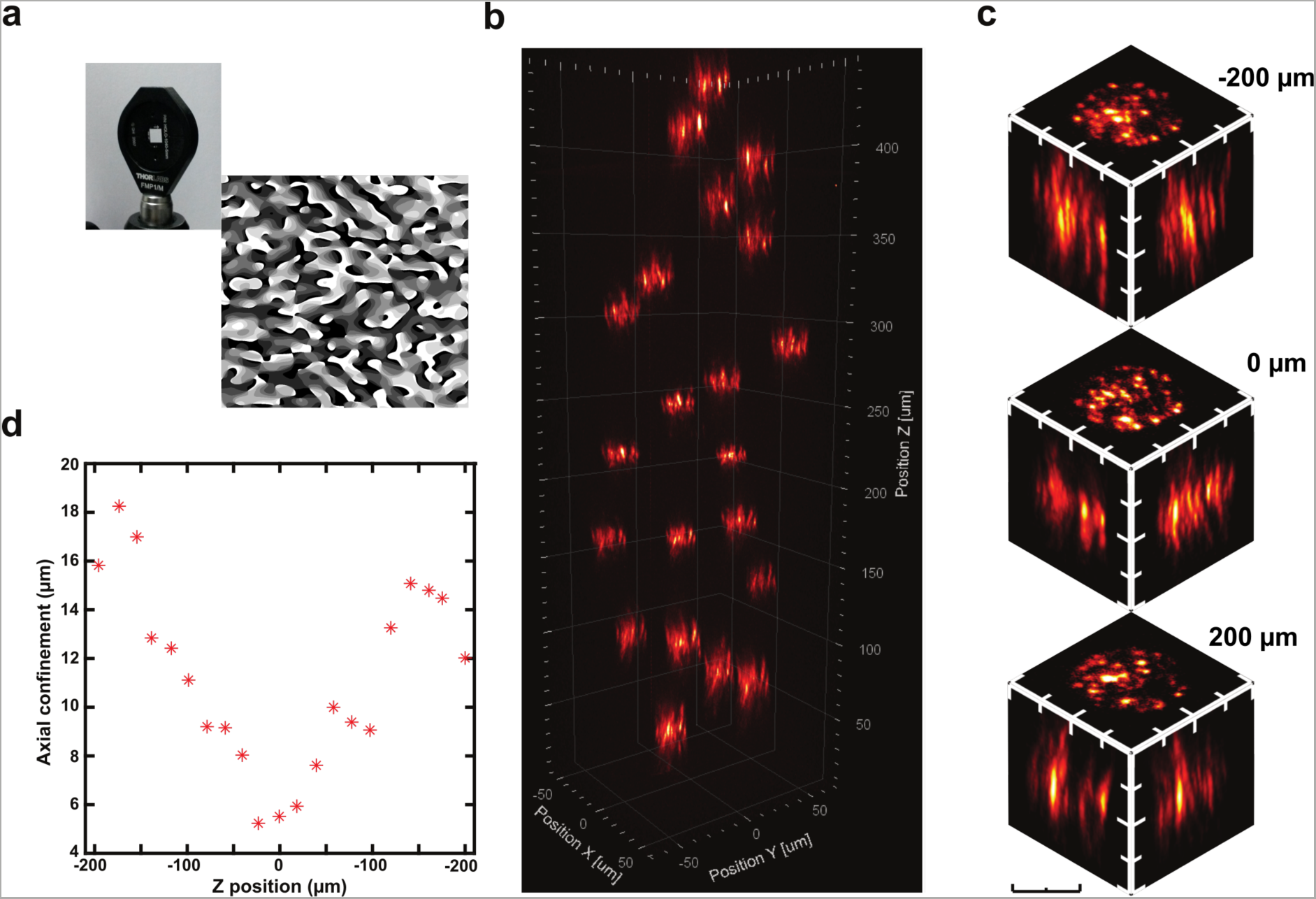
MTF-CGH with a static phase-mask. **(a)** Left: Photo of the holographic static phase mask mounted on a 1-inch circular holder. Right: 8-level computer-generated hologram used to fabricate the static phase mask. (**b**) 2PE fluorescence volume representation of 20 holographic circular spots of 20-μm diameter, each of them lying on a different plane, in a volume of 130×130×400 μm^3^. To record the stack from which we reconstructed the 3D volume, we used 200 mW average laser power at the back aperture of the objective (60x, 0.9 NA), by integrating 1 s per plane with the imaging camera. (**c**) *x*-*y*, *x*-*z* and *y*-*z* projections of three spots, located at *z*=-200, 0, 200 μm from the focal plane. Scale bar: 20 μm. (**d**) Axial confinement, calculated as the FWHM of the axial intensity profile of each spot, as a function of the *z* position. The mean value across the whole FOE was 11.0±4.0 µm FWHM.

Overall, these results demonstrate that our method can produce multiple spatiotemporally focused spots with a <15 μm axial resolution and uniform (within 40% from the average) light distribution across at least 300×300×500 µm^3^. For comparison we show in **Supplementary Fig. 4** the same spot distribution without TF revealing a 3 times larger axial resolution.

**Figure 4:**
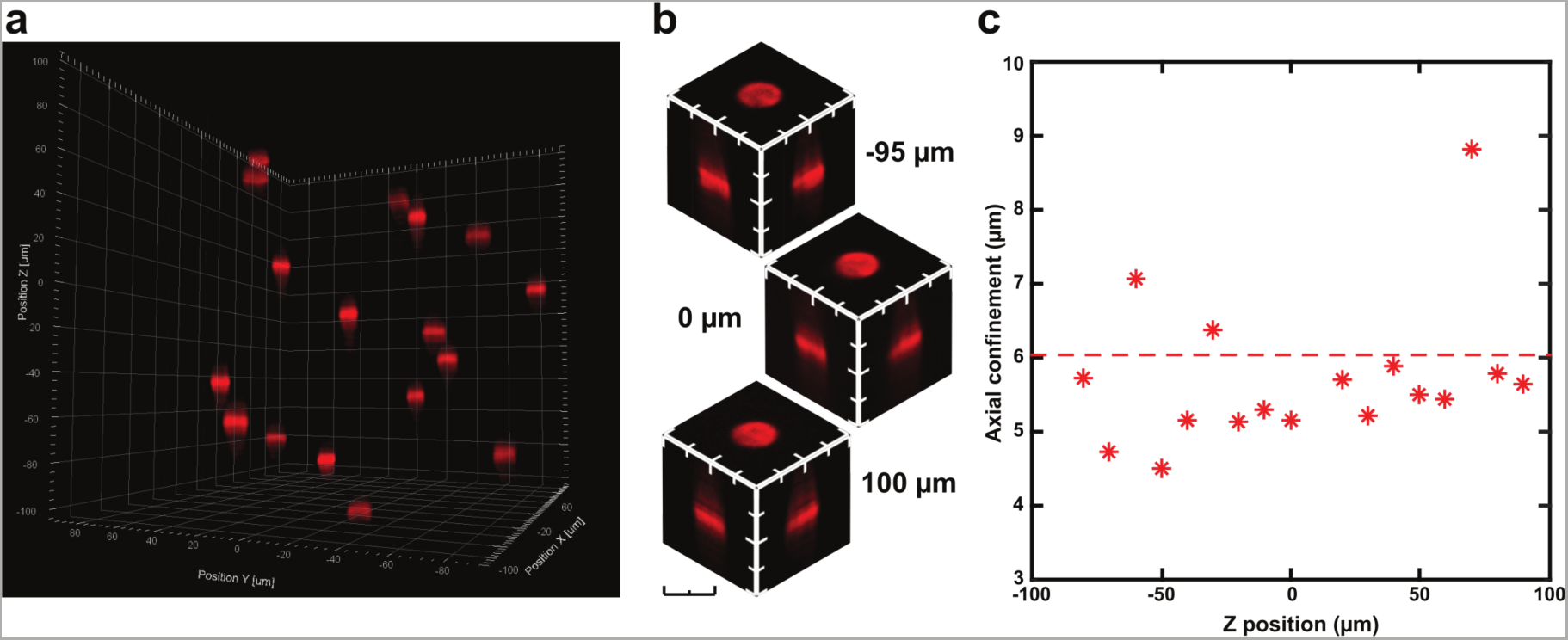
MTF-GPC. (**a**) 2PE fluorescence volume representation of 17 GPC circular spots of 12-μm diameter, each of them lying on a different plane, in a volume of 200×200×200 μm^3^. To record the stack from which we reconstructed the 3D volume, we used 150 mW average laser power at the back aperture of the objective (40x, 0.8 NA), by integrating 2 s per plane with the imaging camera. (**b**) *x*-*y*, *x*-*z* and *y*-*z* projections of three spots, located at *z*=-95, 0, 100 μm from the focal plane. Scale bar: 12 μm. (**c**) Axial confinement, calculated as the FWHM of the axial intensity profile of each spot, as a function of the *z* position. The mean value across the whole FOE was 6.0±1.5 µm FWHM.

### Multiplexed Temporally Focused Computer-Generated Holography with a static phase mask

Next, we replaced SLM1 with a static custom-made 8-grey level phase-mask, which was fabricated to create a 20-μm diameter circular holographic spot (See Methods, **Fig.3a**) when coupled to a 60x objective (NA=0.9). **Fig. 3b** shows the 3D reconstruction of the 2PE fluorescence generated by 21 excitation spots arranged in a FOE of 130×130×400 µm^3^. **Fig. 3c** illustrates the details of three replicas for three different *z* planes (-200, 0, 200 μm), demonstrating the preservation of the spot quality in a *z* range of 400 μm. **Fig. 3d** shows the dependence of the axial confinement for 21 spots in *z*-direction. The mean axial resolution was 11.0±4.0 µm, reaching a minimum value of ~5 µm at the centre of the FOE. Using a larger NA and higher magnification objective improved the axial resolution at the centre of the FOE but also induced a stronger dependence on the axial position (with the 40x objective the FWHM increased by ~0.4% per µm; with the 60x objective it increased by ~1% per μm), since in this case one needs a more divergent beam at the back aperture to obtain the same axial shift, resulting in larger aberrations.

Overall, **Fig. 3b** demonstrates that the generation of multiple spatiotemporally focused spots can be achieved by replacing a dynamic liquid-crystal matrix with a fixed phase mask. This limits the generation of patterns to pre-defined 2D shapes but enables to considerably reduce the complexity (and the total cost) of the optical system.

### Multiplexed Temporally Focused Generalized phase contrast (MTF-GPC)

In a third approach, we changed the beam-shaping unit to a GPC-interferometer (**Fig. 1a**, inset), whose image plane coincided with the grating for TF^34,35^. As **Fig. 4** demonstrates, by coupling such a system with the holographic multiplexing of SLM2, we generated multiplexed temporally focused generalized phase contrast (MTF-GPC) spots on a FOE of 200×200×200 µm^3^. **Fig. 4b** shows the excitation spots at three different planes (-95, 0 and 100 μm) from which one can clearly recognize the uniform speckle-free intensity distribution typical of GPC. In agreement with previous findings^34^, the flatter optical wavefront of GPC enabled achieving a higher axial resolution (6.0±1.5 µm FWHM, (**Fig. 4c**) than the one achievable with CGH using the same objective (**Fig. 2c**). However, as the scheme of **Fig.1a** illustrates, in this configuration the dispersed beam after the diffraction grating was focused on SLM2, resulting in the illumination of only a small portion of SLM2 in the direction perpendicular to TF (*y*-direction, vertical in the optical table). In the TF direction instead, dispersion of spectral frequencies enabled illumination of the full SLM2 chip. Quantitatively, with the optical components used, a spot size of 12-μm diameter at the sample plane determined a vertical beam size at SLM2 of ~1.5 mm. To prevent power induced stress of the liquid-crystal array, we limited the average laser power that illuminated SLM2 to ~ 0.5 W, which in turns limited the total number of spots that we could project to roughly 20.

These results demonstrate that MTF-GPC enables the generation of multiple speckle-free spatiotemporally focused spots with optimal axial resolution. For applications requiring generation of several tens of spots, solutions to enlarge the illumination of SLM2 along the vertical direction are necessary.

## In-vivo high resolution multi-cell targeting

We finally applied our reference technique for the 3D generation of multiple temporally focused spots, namely MTF-CGH, to two different biological paradigms: the 2P-photoconversion of Kaede protein^7^ in zebrafish larvae (**Fig. 5**) and the 2P photoactivation of superfolder GCaMP (PA-GCaMP) in drosophila larvae^6^ (**Fig. 6**).

**Figure 5:**
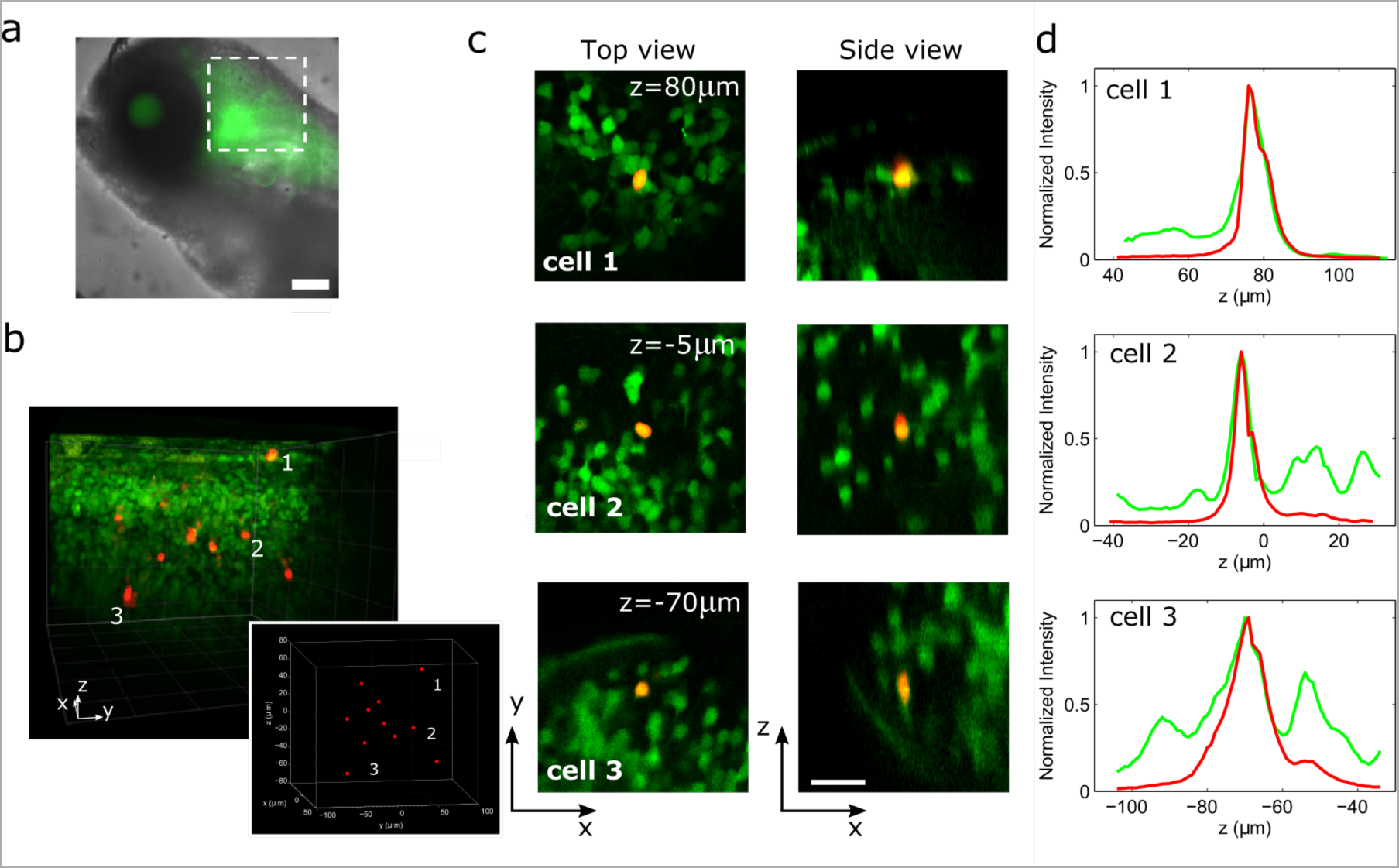
Simultaneous photoconversion of Kaede expressing neurons in zebrafish. (**a**) Superposed brightfield and widefield fluorescence images of the head of a double transgenic *Tg(HuC:gal4; UAS:kaede)* zebrafish larvae. The dashed square represents the approximate area where we performed photoconversion. Scale bar: 100 μm. (**b**) 3D view of a 2P stack (λimaging=780 nm) merging green and red fluorescence after targeted simultaneous 2P photoconversion (λconversion=800 nm) of a set of 11 neurons. Represented volume: 178×178×251 μm^3^. The inset represents the 3D MTF-CGH illumination pattern composed of multiple 6-μm diameter spots used for photoconversion. (**c**) Top and side single frame view extracted from the 2P stack reported in **b**, zooming on 3 representative photoconverted cells (labeled 1-3 in panel **b**). Scale bar: 20 μm. (**d**) Normalized axial intensity profiles of green and red fluorescence integrated over *z* for the 3 cells reported in **c**.

**Figure 6:**
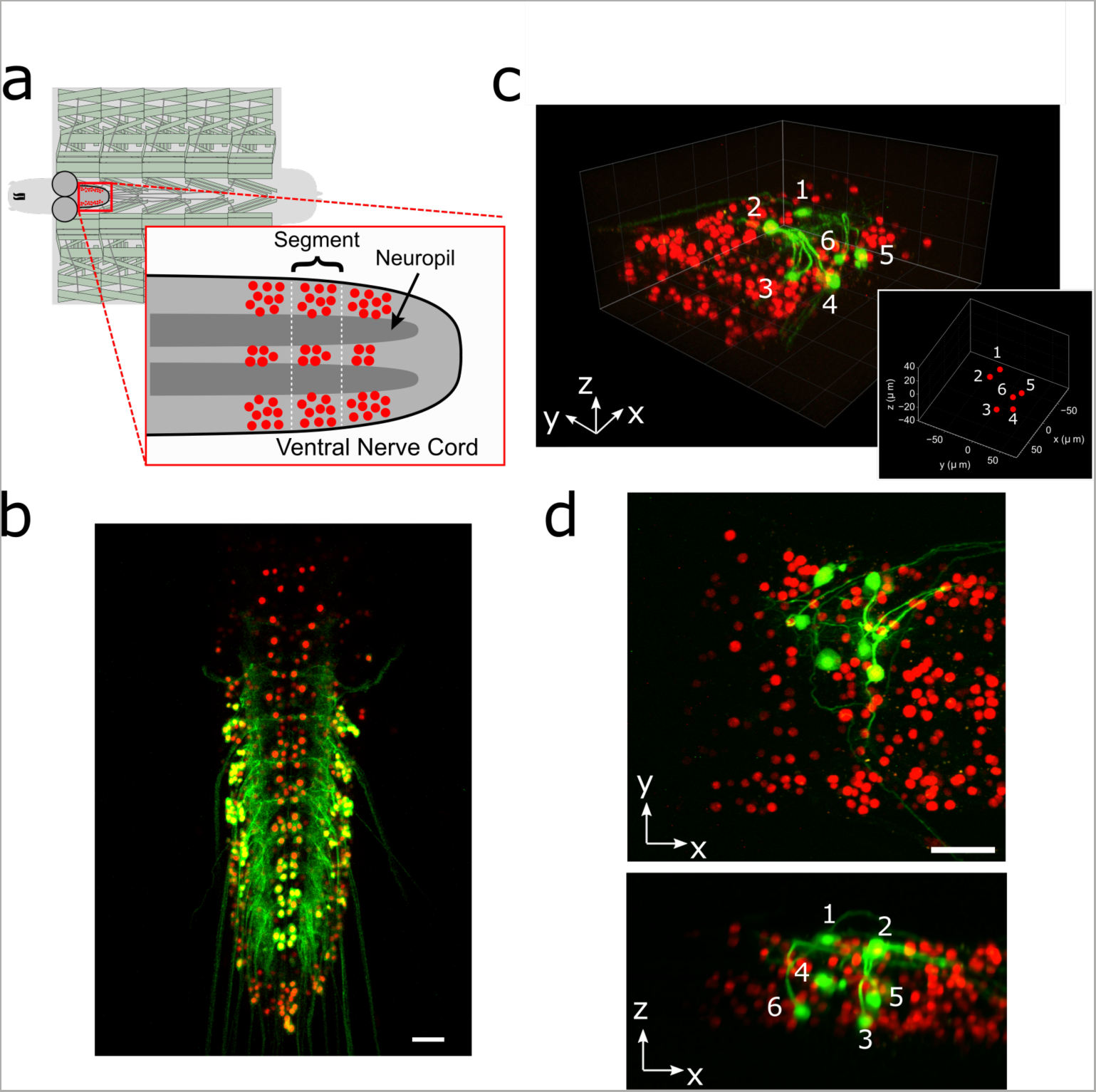
Photoactivation of sPA-GCaMP in drosophila larvae. (**a**) Schematic representation of a dissected drosophila larva. The dissection exposes the ventral cord (red rectangle) in which motor neurons co-express nuclear mCherry (red dots) and photoactivable sPA-GCaMP6f. (**b**) Max projection of a *z*-stack of green (sPA-GCaMP6f) and red (mCherry) fluorescence performed after wide 1P (405 nm) photoactivation of motorneurons of the ventral central cord (see Methods). Image acquired on a confocal microscope. Scale bar: 30 μm. (**c**) 3D view of a 2P stack (λimaging=920 nm) merging green and red fluorescence after 2P (λactivation=760 nm) targeted simultaneous photoactivation of a set of 6 motor neurons (labeled with numbers). Represented volume: 178×178×140 μm^3^. The inset represents the 3D MTF-CGH illumination pattern composed of multiple 5-μm diameter spots used for photoactivation. (**d**) Top (up) and side (down) max projection of green and red fluorescent after photoconversion, corresponding to panel **c**. Numbers label targeted photoactivated neurons. Scale bar: 30 μm.

### Kaede photoconversion in the hindbrain of zebrafish larvae

We prepared samples of zebrafish larvae expressing the Kaede protein in a very dense population of neurons of the hindbrain using *Tg(HuC:gal4;UAS:Kaede)* transgenic larvae (**Fig. 5a**). Kaede is a photoconvertible protein that emits initially green fluorescence and red-shifts its emission under UV^7^ or near-infrared 2P illumination^57^. In order to precisely define and monitor photoconversion patterns, we implemented our MTF-CGH technique in a commercial up-right Zeiss microscope coupled with 2P galvo-based scanning imaging (see Methods). We first acquired a 2P scanning *z*-stack of the green fluorescence from a ~200×200×300 µm^3^ volume in the larva hindbrain (**Fig. 5b**), from which we selected 11 individual neurons, distributed at 11 different depths, for photoconversion. We then precisely tailored the 3D patterned illumination to simultaneously photoconvert these neurons using 2P excitation (λ=800 nm). For this experiment SLM1 generated a circular spot of 6 µm in diameter to excite individual neurons, while SLM2 multiplied such a shape, temporally focused by the diffraction grating, at 11 distinct positions. SLM2 was also used to adjust the relative intensity of each spot to compensate for both diffraction efficiency and losses due to scattering through different depth of the tissue (see Methods).

Finally, we acquired a second 2P scanning *z*-stack to measure both the green and red fluorescence. Photoconversion increased the red fluorescence in the target cells by more than a factor of 15 (19±10 fold; n=11 targeted cells) with respect to neighbouring cells (**Fig. 5b-d**). Top and lateral views from 3 photoconverted cells are reported in **Fig. 5c**. The corresponding normalized fluorescence axial profiles (**Fig. 5d**) show red fluorescence induced solely in the targeted neurons. This confirms the precise targeting and single-cell resolution (FWHM of red fluorescence axial profiles = 7.7 ± 3.3; n=11 cells) of the patterned illumination, down to ~200 μm deep in the brain tissue with minimal photoconversion induced in neighbouring cells, despite the highly packed neuron ensemble.

### Targeted Simultaneous 3D Photoactivation of sPA-GCaMP6f in Drosophilae larvae

We subsequently tested the potential of our system in the central nervous system of *Drosophila* larvae, whose cells expressed a recently developed photoactivable genetically encoded calcium indicator, sPA-GCaMP6f. Such indicator switches from an original dark state to a bright state via UV or 2P infrared illumination^6^.

Larvae were dissected to expose the ventral nerve cord where sPA-GCaMP6f was expressed in all motor neurons (*OK6-Gal4*). These neurons co-expressed a nuclear mCherry (**Fig.6 a,b**). A 2P image of the mCherry fluorescence allowed us to reconstruct the 3D distribution of the sPA-GCaMP6f expressing motorneurons (**Fig. 6a**). We then selected for 2P photoactivation at 760 nm with our MTF-CGH system a subset of 6 individual motorneurons, belonging to a stereotyped group that all projected to a common hemisegment in the larval body (**Fig. 6c**). We generated 6 different 5 µm holographic spots aimed at the cell nuclei. Subsequent imaging of the green fluorescence from sPA-GCaMP6f monitored the photoactivation of sPA-GCaMP6f molecules. As **Fig. 6c-d** shows, green fluorescence increased >15 times only in the targeted neurons. Photoactivation of untargeted neighbouring cells was minimal, despite the very dense distribution of sPA-GCaMP6f expressing neurons and the even denser neuropil containing the processes of the expressing neurons (see **Fig. 6b**, where photoactivation was done by wide-field 1P illumination). Within minutes after photoactivation, neuronal processes of the targeted cells could be clearly distinguished from background (**Fig. 6c-d),** making it possible to track neuronal morphology precisely.

## Discussion

In this work we demonstrated, thoroughly characterized and applied to biological proofs-of-principle a novel optical system enabling the generation of multiple temporally focused illumination targets at arbitrary 3D locations. The system was based on two independent beam-shaping steps: the first one defined the lateral shape of the illumination spot and projected it on the TF grating; the second multiplexed the original shape at the desired positions through CGH.

A unique feature of this configuration is that the total number of excitation spots, as well as the size of the achievable FOE at the target volume are decoupled from the first beam-shaping unit, and only depend on the performances of SLM2 (total number of illuminated pixels, pixel size, number of grey levels). We demonstrated that this characteristic makes our system compatible with several different beam-shaping approaches such as dynamic CGH, CGH with a static phase mask and GPC interferometry.

Among the approaches that we tested, dynamic CGH, which uses a reconfigurable liquid crystal SLM, enables maximal flexibility and quicker lateral shaping. Replacing the bulky SLM with a smaller static phase mask reduces the flexibility of the system but leads to a simpler and more compact optical design. Moreover, as the first beam-shaping units in many cases only needs to generate simple 2D shapes (round spots), one can choose a low-resolution (e.g. 8-grey-levels) cost-effective phase mask. A phase mask made up of a single plate with different holograms imprinted on it will enable to easily change the spot size and shape and hence to cover more applications.

Similar to dynamic CGH, GPC enables a quick adjustment of the spot size and shape. Moreover, it generates illumination patterns with superior axial resolution and highly uniform (speckle-free), which is particularly advantageous over CGH for applications requiring spots sizes comparable to the speckle size. In the past, the main limitations of GPC were the reduced FOE imposed by the constrain of keeping the ratio of the illuminated over the non-illuminated area in the FOE to 0.25^34^, as well as the lack of 3D light shaping capability. Using SLM2 for axial and lateral multiplexing removes both these constrains, which makes MTF-GPC a very promising approach. Yet, the <2 mm vertical size of the beam at SLM2 limits the power that can be used and therefore the maximum number of achievable targets. At the same time, using only a limited number of pixels in the vertical direction limits the performances of SLM2 and deteriorates the quality of the generated holograms. Adding an Echelle grating to the TF setup to fill the SLM2 area by using diffraction in one dimension and spectral dispersion in the other^58^, or cylindrical lenses to independently tune the beam size in the *x* and *y*-directions should allow removing these limitations.

In all cases, the total FOE at the sample position depends on the SLM2 pixel size and the telescope used to conjugate the SLM2 plane with the objective back aperture. When using the 40x, 0.8 NA objective our system had a theoretical FOE^13^ of 750×750×990 µm^3^. Experimentally we could demonstrate <15 μm axial resolution and uniform intensity within 40% from the average within a FOE of 300×300×500 µm^3^, limited by the size of the optics (mirrors and lenses) placed after SLM2. An optical design using larger optical elements and careful aberration correction should enable to reach the theoretical FOE.

Very recently, we demonstrated a first implementation of 3D CGH with TF using two SLMs, with the advantage of allowing different shapes to be projected at different axial positions^13^. However, that approach required tiling of the two SLMs in multiple holograms, therefore compromising the spot quality when increasing the number of planes and imposing a physical limit to the total number of addressable planes. The alignment of the system was complicated as the method also required a perfect match between the tiles of the two different SLMs and a more complex calibration procedure to generate targets with uniform intensity was necessary. Finally, as the lateral and axial displacement of the 3D spots was performed by different SLMs, the performances of the systems (number of spots and FOE) were also determined by the properties of SLM1. This compromised the compatibility of that approach with the use of a fixed phase mask or GPC. All these limitations are overcome with our novel method.

A similar configuration to the one described here, using as first beam-shaping unit an expanded Gaussian beam, a grating for TF and one SLM for multiplexing, recently permitted the 3D generation of 2.5-μm-diameter spatiotemporally focused spots, with axial resolution of 7.5 μm^51^. Such method is however limited to Gaussian shapes and does not enable easy spot-size reconfiguration. Moreover, similar to the case of GPC, the use of expanded Gaussian beams under-fills SLM2 in the non-TF direction. Adesnik and colleagues^59^ could overcome this issue by placing SLM2 in a position at which the beam was not focused. As a downside however, the spatial and temporal foci at the sample plane did not coincide anymore, thus significantly broadening the achievable axial resolution. Finally, both approaches used temporally focused Gaussian excitation shapes, which in contrast to the use of top-hat intensity profiles, typical of CGH and GPC, experience a power-dependent lateral resolution and worse propagation through scattering media^35^.

We applied the MTF-CGH system to the *in vivo* 2P conversion of Kaede protein in the brain of zebrafish larvae and to the *in vivo* 2P activation of a photoactivable version of GCaMP (sPA-GCaMP6f) in the central nervous system of fruit flies. In both applications, we demonstrated in-depth simultaneous targeting of multiple individual neurons. Temporally focused patterns, as already demonstrated in previous works^35,13^, are robust against propagation through scattering media, which allowed individual neurons within a highly packed ensemble to be precisely targeted up to a depth of 200 µm, with minimal spurious fluorescence induced in neighbouring cells. The primary limitation in photo-converting a higher number of targets was the total laser power available at each position. Longer exposure times combined with the introduction of real-time movement correction (as well as the use of more powerful lasers) will allow increasing the number of targeted neurons.

The capability of the optical system to precisely target multiple cells can be applied to a large variety of photo-switchable proteins^8,60^, and could be useful for tracking specific cellular ensembles *in vivo* and during development. In particular, targeted photoactivation of calcium indicators opens the way to the simultaneous morphological and functional investigation of specific neuronal sub-circuits where cells are either too dense for traditional analysis, or where there is a lack of cell-specific genetic driver lines^6,15^.

Importantly, MTF light shaping can be combined with optogenetics to precisely manipulate neuronal circuits in mm^3^ volumes. We recently demonstrated that the combination of efficient opsins, such as Chronos, ReachR and CoChR, with holographic stimulation using an amplified fibre laser, leads to the generation of APs with unprecedented temporal resolution and precision both *in vitro*^48,47,46^ and *in vivo*^61^. In that work we generated APs with <3 ms temporal resolution and <0.1 ms temporal jitter, using excitation spots with diameter of 10-15 µm in diameter and power densities in the range of 0.03-0.1-mWμm^−2^. We can therefore estimate that, especially when using last generation high-power lasers (delivering tens of Watts), MTF light-shaping could easily lead to the simultaneous activation of hundreds of cells. Under such illumination conditions, it is very important to keep the amount of heat generated in the tissue within a certain safety range. However, we recently demonstrated that, as long as the multiple illumination targets are placed at an average distance larger than the corresponding heat diffusion length (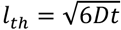, with D thermal diffusivity, and t illumination time), the total temperature rise induced in a brain tissue is comparable to the case of a single illuminated target, which, for 3 ms pulse duration and 0.1 mWμm^−2^ power density, does not exceed 1°C^62^.

The demonstrated MTF light shaping technique could therefore constitute the basis of a reliable and robust approach for 3D ‘all-optical’ brain circuits control on large scales, especially if combined with 3D imaging techniques^22,63–65^. At the same time, its use is not only limited to neuronal activation, but could as well be extended, in combination with camera detections^24,25^ and multi-plane spatial demixing algorithms^66^, to fast volumetric functional imaging, using calcium or voltage sensors. More generally, any application relying on light patterning methods and non-linear phenomena, such as photo-polymerization^67^, optical data storage and photolithography^68^, will benefit from this technique.

## Methods

### MTF-CGH setup

The optical system is schematically depicted in **Fig. 1**. The laser source used in the experiment was a femtosecond fibre laser (Fidelity 18, Coherent), emitting at 1040 nm, delivering 140 fs pulses at a pulse repetition rate of 80 MHz, with an average power of 18 W. The laser beam was expanded 10 times to fit the active area of SLM1 (LCOS-SLM X10468-07, Hamamatsu Photonics, resolution 800×600 pixels, 20 μm pixel size), which modulated the phase of the incoming beam through a standard Gerchberg-Saxton algorithm^69^ to create the desired 2D intensity pattern. The image of the pattern was formed through the lens L1 (f1=500 mm) on a blazed reflectance diffraction grating (830 l/mm, 53004ZD02-035R, Richardson Gratings; G) for TF. The grating was aligned such as the first diffraction-order was diffracted along the optical axis of the microscope, perpendicular to the grating plane. It was subsequently collimated by lens L2 (f2=500 mm) and impinged on SLM2 (LCOS-SLM, X13138-07, Hamamatsu Photonics, 1280×1024 pixels, 12.5 μm pixel size), imaged at the back focal plane of the excitation objective (Olympus LUMPLFL40xW/IR2, NA 0.80 objective; OBJ) via lenses L3, f3=1000 mm and L4, f4=500 mm). The phase modulation created by SLM2 produced a set of 3D diffraction-limited spots, which multiplexed the original 2D pattern. For precise axial displacement, the phase profile applied to SLM2 was calculated with a weighted Gerchberg-Saxton algorithm^55^, modified to include the non-parabolic terms in the description of the microscope objective^41^.

Zero order, i.e. the part of light that is not modulated by the SLMs, was physically blocked for both SLMs. This created an inaccessible region in the central part of the FOE. To make use of the entire FOE, other solutions for suppressing the excitation effect of the zero order could be considered, like adding one or a pair of cylindrical lenses in front of the SLMs^70^, or adding a destructively interfering spot to the phase hologram design^71,72^.

For characterising the performances of the system, 2PE fluorescence from a thin (~1 μm) spin-coated fluorescent layer of rhodamine-6G in polymethyl methacrylate 2% w/v in chloroform was collected by a second microscope objective (Olympus UPLSAPO60XW, NA 1.2) in transmission geometry and detected with a CCD camera (CoolSNAP HQ2, Roper Scientific), using a dichroic mirror and a short pass filter for rejecting laser light (Chroma Technology 640DCSPXR; Semrock, Brightline Multiphoton Filter 750/SP). For 3D reconstruction of illumination volumes, the ‘imaging’ objective imaged the rhodamine-layer plane throughout all the experiment, while the excitation objective OBJ1 was scanned over the desired *z* range with a piezoelectric scanner (PI N-725.2A PIFOC). The CCD camera and the piezoelectric scanner were controlled by LabVIEW. A custom-designed software, Wavefront-Designer IV^73^, written in C++ and using the open graphic library Qt 4.8.7, controlled the two SLMs and run the Gerchberg-Saxton-based algorithms. The software also includes the phase corrections for the first orders Zernike aberrations.

We performed the analysis of the recorded stacks with Matlab, ImageJ and the Imaris software (Bitplane, Oxford Instruments).

The 2PE fluorescence values for each spot were obtained by integrating the intensity of all the pixels in a circular area containing the spot, in the plane where the intensity was at its maximum value (i.e. the TF plane). Axial intensity distributions were obtained by integrating the intensity of the pixels in the same area for each plane of the recorded stack. Reported values for the axial confinement were the fit of the axial profile of the spots with a Lorentzian model and referred to the FWHM of the curves. Statistical data in axial resolution measurements were reported as mean±standard deviation.

The static phase mask, used alternatively to SLM1, was fabricated by photolithography (Double Helix Optics, LLC) on the base of an 8-grey-level phase profile calculated with the Gerchberg-Saxton algorithm (**Fig. 3a**). The dimensions of the encoded area of the mask were 5×5 mm^2^, thus beam expansion was adjusted in that case to 5x, to fit the size of the phase mask. A 2:1 telescope (f1PM=200 mm, f2PM=400 mm) was used to magnify the mask approximately to the size of SLM2, and generated a 20-μm diameter circular spot at the focal plane of the optical system using a 60x, 0.9 NA objective (Olympus LUMPLFL60xW/IR2).

### MTF-GPC setup

The beam shaper for GPC patterns is schematically shown in the bottom inset of **Fig. 1a**. Similarly to the setup previously described^34^, SLM1 (LCOS-SLM; Hamamatsu Photonics X10468-07, resolution 800×600 pixels, 20 μm pixel size) was illuminated at oblique incidence by the expanded laser beam (10x). The device was controlled by a version of Wavefront-Designer IV that, given a target intensity distribution at the focal plane of the microscope objective, converted the intensity map into a binary phase map and applied the output profile to the SLM. A first achromatic lens (L1GPC, f1GPC=300 mm) focused the reflected beam from the SLM on the phase contrast filter (PCF), positioned at the Fourier plane of the lens. For optimal phase contrast, the round-PCF pitch size was chosen in the range of 80-100 μm diameter, and phase-shifted by ~λ/2 the low-spatial-frequency components^74^. A second achromatic ^lens (L2GPC, f2GPC=60 mm) recombined the high (signal wave) and low (synthetic reference wave) spatial^ frequency components^75^. The introduced phase-shift caused those two components to interfere and produced an intensity distribution, according to the spatial phase information carried by the higher spatial frequencies, at the grating plane for TF. For GPC, we replaced the diffraction grating with one of 1200 lines/mm (Optometrics Corporation, G1200R750JGAS), in order to fill SLM2 in the TF direction. Due to the focalization of the beam, in the direction perpendicular to TF (*y*-direction, vertical in the optical table) the illuminated region of the SLM2 remained smaller (~1.5 mm). To prevent any possible damage to the SLM liquid crystal, we limited the total delivered average power on SLM2 to 0.5 W. This value was chosen in accordance with the characterization performed by the manufacturer (Hamamatsu Photonics) that did not observe, at similar average power density and at even higher peak powers, any damage or phase modulation changes after multiple hours exposure.

The setup after the grating was the same as the one described for CGH. Similarly, the same procedure as described above was followed for collecting and analysing fluorescence data.

### Optical system used in biological experiments

The biological experiments presented here were performed in a microscope using MTF-CGH with two SLMs for photostimulation and a commercial 2P scanning microscope with galvanometric mirrors (VIVO, 2-PHOTON, 3i-Intelligent Imaging Innovations). The whole setup was built around a commercial upright microscope (Zeiss, Axio Examiner.Z1). The laser used in this case was a Ti:Sapphire oscillator (pulse width ~100 fs, repetition rate 80 MHz, tuning range 690-1040nm, Mai Tai DeepSee, Spectra-Physics). The photostimulation path consisted of a lens system (fBE1=19 mm and fBE2=150 mm) which expanded the laser beam ~8 times before SLM1 (LCOS-SLM X10468-07, Hamamatsu Photonics, resolution 800×600 pixels, 20 μm pixel size). The intermediate holographic images were then focused by a f1=500 mm lens on the diffraction grating (830 lines/mm, Item No. 55262, Edmund Optics), followed by a second lens, f2=500 mm, the SLM2 (LCOS-SLM X13138-07, Hamamatsu Photonics, 1280×1024 pixels, 12.5 μm pixel size) a 4-f telescope (f4= f5 =500 mm), and, finally, the last telescope before the objective with f6=300 mm, f7=250 mm. The objective used for the experiments was a water immersion 40x-NA 0.8 objective LUM PLAN FI/IR, Olympus.

The holographic path was then coupled with the commercial 2P scanning module, made of 2P galvo-scanner of (2P Vector, 3i) and a detection based on two spectrally resolved GaAsP photomultipliers. Acquisition was controlled by SlideBook6 software (3i). The two illumination paths were recombined on a polarizer cube and shared the f4=250 mm lens before entering the microscope. The switch between the optical path for scanning imaging and holographic illumination was performed thanks to a movable mirror driven by a servomotor. Illumination power and pulses were controlled by a Pockels cell (350-80, Conoptics).

### Kaede photoconversion

*Zebrafish housing and handling:* All procedures were approved by the Institut du Cerveau et de la Moelle épinière (ICM) and the National Ethics Committee (Comité National de Réflexion Ethique sur l’Expérimentation Animale Ce5/2011/056) based on E.U. legislation. Embryos were raised in an incubator at 28.5°C under a 14/10 light/dark cycle until the start of experimentation. Experiments were performed at room temperature (23-26°C) on 2-6 days post fertilization (dpf) larvae. All experiments were performed on Danio rerio embryos of AB and TL background and mitfa -/- animals were used to remove pigments above the hindbrain. For photoconversion experiments, we used double transgenic *Tg(HuC:gal4; UAS:kaede)* larvae where the HuC promoter drives pan-neuronal expression of Kaede at the larval stage. Embryos were dechorionated and screened for green fluorescence at 1 dpf. Larvae screened for Kaede fluorescence were later embedded laterally in 1.5% agarose. Larvae were anaesthetized in 0.02% tricaine (MS-222, Sigma-Aldrich, USA).

#### Photoconversion and imaging

A first 2P imaging *z*-stack of green fluorescence, was recorded with the scanning laser at 920 nm to map the location of neuronal cells. By using a Matlab-based graphic interface, we selected a subset of neurons for photoconversion and defined accordingly a multispot pattern made of circular spots of 6 μm diameter at 800 nm illumination. As the zero order from SLM2 consisted also of a circular spot that was identical to the multiplexed replica, we photoconverted an additional neuron located at the centre of the FOE. The corresponding holograms for SLM1 and SLM2 were calculated by using our custom-designed software, Wavefront-Designer IV^73^. The relative intensity between spots was adjusted according to their position in order to compensate for both diffraction efficiency and tissue scattering (low diffraction efficiency or deeper regions) by sending higher intensities to positions where spots were dimmer. This compensation was obtained by adjusting the relative weights of each spot as input of the iterative algorithm for phase calculation of the hologram in SLM2^213,54,55^. To minimize thermal damage during photoconversion, we delivered trains of 10-ms pulses at 50 Hz with total laser power around 130 mW (corresponding to an average illumination density of ~0.4 mW/μm^2^, see **Fig. 5**), for periods of time that ranged from few tens of seconds to four minutes.

After photoconversion, we acquired a second *z*-stack in the green and red channels with the scanning laser tuned to 780 nm in order to efficiently excite red-photoconverted fluorescence and minimize crosstalk of green fluorescence in the red channel. The use of low laser power and multiple frame averages minimized artefactual photoconversion induced by the scanning laser. Increase in red fluorescence in target cells was estimated by comparing their red emission after photoconversion with the average red emission from five non-targeted coplanar neighbouring cells that were randomly selected. Axial intensity profiles corresponding to **Fig. 5c,d** were obtained by integrating red and green fluorescence intensity over an ROI covering the targeted cell body with axial correction for a *z*-shift below the targeting stack axial sampling rate.

### sPA-GCaMP photoactivation

#### Drosophila preparation

We used female wandering third instar *Drosophila* larvae, expressing sPA-GCaMP6f and mCherry-nls in all motor neurons under the control of OK6-Gal4^76^ with the following genotype *w*^1118^;OK6-Gal4/UAS-mCherry-nls;UAS-sPA-GCaMP6f/UAS-sPA-GCaMP6f). Larvae were ^dissected in HL3 saline containing, in mM: 70 NaCl, 5 KCl, 0.45 CaCl2∙2H2O, 20 MgCl2∙6H2O, 10 NaHCO3, 5^ trehalose, 115 sucrose, 5 HEPES (pH adjusted to 7.2). The CNS and peripheral motor neurons were exposed by making a longitudinal dorsal incision, removing all organs, and pinning the cuticle flat.

*Photoconversion and imaging:* The experimental protocol was similar to what performed for Kaede photoconversion experiments. The initial reference ventral nerve cord 2P stack was performed at 920 nm, to record nuclear mCherry fluorescence. Photoactivation was performed with a 760 nm patterned illumination made of multiple circular spot of 5 µm diameter. We used trains of 100-ms pulses at 5 Hz with total laser power around 130 mW (corresponding, in the case of **Fig. 6**, to an average illumination density of ~1 mW/um^2^) and for periods of time that ranged from 1 up to 4 minutes. After activation, a 2P *z*-stack with the scanning laser at 920 nm was acquired to reconstruct both mCherry and activated sPA-GCaMP6f fluorescence. As a reference for total motor neuron density and sPA-GCaMP6f expression, large field 1P photoactivation (shown in **Fig. 6b**) was achieved separately (Zeiss LSM780 confocal microscope) by scanning the whole ventral nerve cord with a 405 nm diffraction limited spot and imaging with 488 nm and 561 nm excitation.

## Acknowledgments

We thank Vincent de Sars for software developing, Leslie Kimerling (Double Helix LLC) for providing the phase mask, Coherent Inc. for the loan of the Fidelity laser, Intelligent Imaging Innovations, Inc. for providing the 2P scanning microscope, Martin Booth for useful discussions, Anna Segú-Cristina for help in the preparation of the zebrafish sample, SCM (Service Commun de Miscroscopie – Faculté des Sciences Fondamentales et Biomédicales - Paris) for providing the software Imaris (Imaris v8.4, Bitplane, software available at <u>www.bitplane.com</u>), Sophie Nunes-Figueiredo from the ICM zebrafish facility for animal care, Sorbonne Universités, UPMC Univ. Paris 06, Inserm, CNRS, AP-HP, Hôpital Pitié-Salpêtrière), the team of Jean-René Huynh (Institut Curie, Paris) and of Bassem Hassan (ICM, Paris) for their help with the Drosophila’s transgenic lines. This research was developed with funding from the Defense Advanced Research Projects Agency (DARPA), Contract No. N66001-17-C-4015. The views, opinions and/or findings expressed are those of the author and should not be interpreted as representing the official views or policies of the Department of Defense or the U.S. Government. It received support as well from the ‘Agence Nationale de la Recherche’ (grant ANR-15-CE19-0001-01, 3DHoloPAc and ANR-14-CE13-0016, Holohub), the Human Frontiers Science Program (Grant RGP0015/2016), Getty Lab, the National Institute of Health (Grant NIH U01NS090501-03). N.A. received funding from the European Union’s Horizon 2020 research and innovation programme under the Marie Skłodowska-Curie grant agreement no. 746173. C.W. received funding from the European Research Council (Starting grant #311673) and the New York Stem Cell Foundation as a Robertson Neuroscience Investigator.

